# A scalable mesh microelectrode array platform for longitudinal electrophysiology in neural spheroids

**DOI:** 10.64898/2026.07.15.738387

**Authors:** Tom Stumpp, Fulya Ersoy, Michael Mierzejewski, Meike Beer, Angelika Stumpf, Lisa-Marie Erlandsdotter, Udo Kraushaar, Peter Loskill, Peter D. Jones

## Abstract

Electrophysiological interfacing remains difficult in three-dimensional in vitro models, when using planar microelectrode arrays or optical methods. This challenge limits experimental progress using such models, despite their promise of better physiological relevance than monolayer cell culture. Mesh MEAs which can integrate conformally on or even within tissue offer a possible solution but are not yet widely accessible. Here, we present a mesh MEA device, designed as a simple, manufacturable platform for spheroid electrophysiology. In neural spheroids, the device enabled longitudinal electrophysiological recordings and pharmacological modulation of spontaneous electrical activity. On native polyimide meshes, spheroids maintained their shape while cells enveloped the mesh, embedding electrodes to a depth of 100 µm after 2 weeks. In contrast, laminin biofunctionalization of the mesh promoted outgrowth and migration of cells. This device and associated methods should be adaptable to organoids, ex vivo tissue, or bioengineered in vitro models.

## Introduction

The challenge of understanding human physiology and disease and the limitations of animal models motivate the development of 3D human in vitro models. Microphysiological systems, including organ-on-a-chip devices and organoids, promise to accelerate understanding of diseases and development of treatments. Limited access to human brain tissue and low relevance of animal brains has motivated the development of neural organoid models.^1^ Disorders of the nervous system are the leading cause of disease burden worldwide.^2^

The electrical activity of neurons is an important feature of the brain, which should be reflected in models.^3^ This activity can be studied from the level of ion channels and synapses, up to microcircuits, brain regions, and finally the innervation of the body. Adherent monolayer cell culture is well established for models studying changes to ion channels and synapses. For this, planar microelectrode arrays (MEAs) are an established tool.^4^ These devices are produced by photolithographic structuring of electrodes, traces and insulators on solid surfaces (commonly glass, silicon, or a polymer). Such electrodes are straightforward to produce on planar surfaces, and are ideally suited for monolayers of adherent cells, with direct contact for measurement or stimulation. This concept works well for passive MEAs with tens to hundreds of electrodes or active CMOS MEAs with thousands of densely packed electrodes.^4^

To study microcircuits and connections between more distant regions, contrasting models are under development. Top-down bioengineering strategies use microfluidics^5–7^ or bioprinting^8^ to arrange different cell types and guide their connections. In contrast, organoids offer a bottom-up approach, starting from stem cells and guiding intrinsic developmental processes. Organoids show promise for investigating microcircuits, while assembloids can model connections between brain regions or to other organs.^9^ The inherent three-dimensional (3D) nature of organoid models limits the suitability of conventional MEA devices. Planar devices interface primarily with superficial cells,^10^ solid devices obstruct medium supply,^11^ and penetrating devices disrupt tissue.^10^

In developing new devices, it’s first important to consider the aims of organoid models. A common goal for electrophysiology is to resolve the activity of as many individual neurons as possible.^12^ An ideal device would non-invasively distribute electrodes throughout tissue and record from all cells with high fidelity. For monolayer cell culture, CMOS MEAs may approach this ideal by sampling an entire network at subcellular resolution.^13^ But in organoids, as in the brain,^12^ we expect that dense electrodes and solid devices cause harm – whether placing an organoid on a device or implanting a device into an organoid. This has inspired many groups to investigate mesh MEAs.

Mesh MEAs consist of fine polymer filaments hosting microelectrodes. Their open structure is minimally disruptive yet stable enough to support 3D tissue integration and long-term experiments. Meshes can conform to the surface of tissue^14^ or be enveloped by cells to bring electrodes into the tissue.^15,16^ The low bending modulus resulting from material properties and geometry of the mesh reduces mechanical mismatch. In vivo mesh MEAs elicit minimal immune response and record stable signals over months.^17^ Implantation however requires novel methods.^18^ Flexible meshes can better integrate with tissue in vitro,^16^ yet handling is more challenging than using classical solid MEAs. This extends even to device fabrication: flexible polymer chips are more difficult to handle than silicon or glass chips.

Despite important progress in mesh-based interfaces for 3D in vitro models,^10^ many reported devices remain technically demanding to fabricate, assemble, or use reproducibly across larger numbers of samples. For routine studies of spheroids and organoids, a practical interface must therefore balance biological integration with experimental robustness,^19^ including straightforward handling, compatibility with established MEA workflows, and scalable device production.

In this context, we present the design, fabrication and validation of a mesh microelectrode array device for 3D tissue models (**Figure 1**). We demonstrate its suitability for recording spontaneous electrical activity within neural spheroids produced from a reference human iPSC line via neural progenitor cells. Activity was recorded longitudinally for up to 2 weeks. Application of pharmacological compounds in the mesh MEA device confirmed the neuronal origin of activity. The device allows non-invasive recording from within intact spheroids, as cells envelop the mesh and its electrodes. We also report laminin-based biofunctionalization of the mesh to promote outgrowth of neurons, suggesting possibilities for engineering of assembloids. Beyond electrical activity, we demonstrate how noise and impedance spectroscopy can monitor cells and tissue on the mesh.

**Figure 1.**
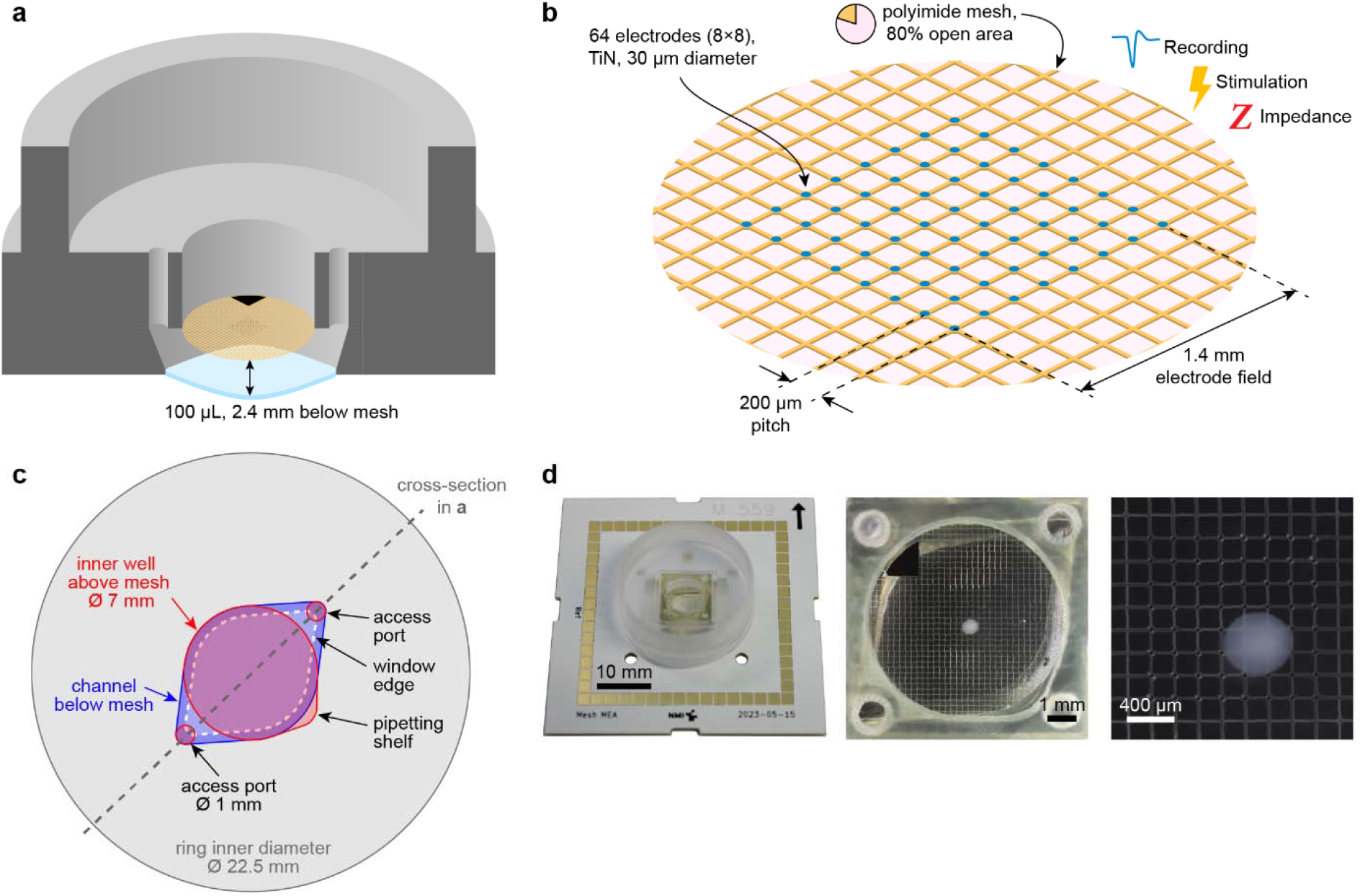
Concept and design of our mesh MEA devices. **a:** Cross-sectional view of the device. Meshes are suspended 2.4 mm above a transparent window. The inner well above the mesh is 4 mm tall. Two ports (1 mm diameter) access the volume below the mesh. The mesh hosts 64 microelectrodes and a triangular reference electrode. **b:** The mesh filaments host 64 microelectrodes at the nodes of a square grid (200 µm pitch). The area of the mesh is 80% open. TiN microelectrodes with a diameter of 30 µm are suitable for recording, stimulation and impedance measurements. **c:** Schematic top view of device. Grey: the outer well. Red: inner well above the mesh, including a pipetting shelf, and 1 mm access ports. Blue: volume below the mesh (lemon-shaped). The volume below the mesh is tapered to the window outline (dashed white line). The diagonal dashed grey line shows the location of the cross-section in **a. d:** Photos of a mesh MEA device and magnifications of a neural spheroid on the mesh.

## Results

### Device design

Our mesh MEA device consists of three parts: the microfabricated mesh having 64 microelectrodes and a reference electrode, a well defining the cell culture volume, and an electrode interface board (EIB) for connection to amplifiers. Rivet bonding connected meshes to the EIB. Wells were assembled with a bottom part below the mesh and a top part above the mesh, and all parts were adhesively bonded (**Figure 2**).

**Figure 2.**
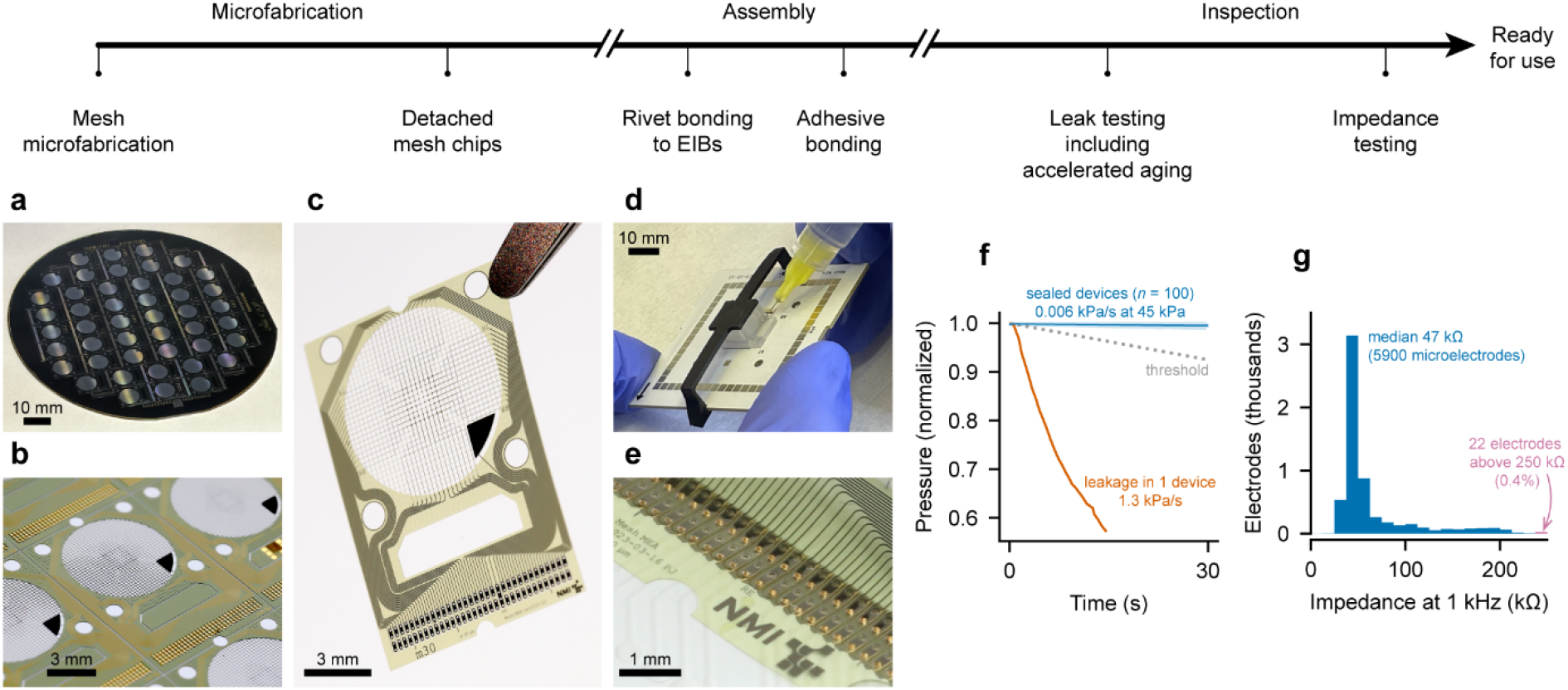
Device fabrication and quality control. Above: flow chart of main microfabrication, assembly, and inspection steps. **a:** 46 mesh chips on a 100 mm carrier wafer. **b:** Macro photo focusing on a single 9×13 mm^2^ chip on the wafer. **c:** A single chip detached from the wafer, held by tweezers. The mesh spans the circular region, with central microelectrodes and a triangular reference electrode. Outside of the mesh are holes with a 1 mm diameter: 2 for alignment during assembly, and 2 for access ports. The lower end of the chip contains bond pads. **d:** Application of adhesive to a partially assembled mesh MEA device. Well parts are clamped on both sides of the mesh between the EIB and the black clip. **e:** Rivet bonds connecting the mesh chip to the EIB. The bond pad of each channel has 2 through-holes for rivet bonding. **f:** Leakage testing of 100 mesh MEA devices vs. one defective device. A discretionary threshold is shown in grey. **g:** Histogram of impedance measurements of 100 mesh MEA devices (5900 microelectrodes). The median impedance magnitude at 1 kHz was 47 kΩ, and 99.6% of electrodes were functional.

The mesh follows a square grid with a pitch of 200 µm over a circular region with a diameter of 7 mm. Electrodes are at grid nodes in a central 8×8 array, and are connected by traces extending along filaments, before being routed to bond pads. Filaments are 21 µm wide so that the mesh is 80% open. Any spheroids with a diameter of 180 µm or larger will remain above the mesh.

Using a wire-bonder, gold rivet bonds connect the electrodes to an EIB.^20^ Here, we have designed a 60-pad EIB suitable for headstages such as those from Multi Channel Systems. We have also developed EIBs for card-edge connectors which allowed interfacing to all 64 microelectrodes and the reference electrode (**Figure S1**). This moved all contact pads to one edge and allowed connection to an amplifier even when permanent perfusion tubes were connected.^21^

The well is assembled from multiple polycarbonate components and an epoxy adhesive. The mesh partitions the well into a volume below the mesh (100 µL) and above the mesh. The mesh is suspended 2.4 mm above a transparent bottom. Above the mesh, a 4 mm-tall inner well has a diameter of 7 mm (volume 160 µL), before expanding to a larger well with an inner diameter of 22.5 mm and height of 5.6 mm (volume 2.2 mL). The outer well diameter of 23.9 mm is compatible with membrane chambers.^22^

Two access ports allow filling or medium exchange below the mesh. In the inner well, we have included a small shelf, designed for pipette placement for medium removal without damaging the mesh.

In our experiments, the inner well below and above the mesh was filled with medium (250 µL). Alternatively, spheroids could be cultured at the air–liquid interface (100 µL) or with higher medium volume in the outer well.

### Device fabrication and quality control

Mesh chips were fabricated on 100 mm silicon carrier wafers before being detached (**Figure 2a–c**). Each chip was wire bonded to an EIB, then well components were assembled around the mesh and joined by adhesive bonding (**Figure 2d–e**). In addition to optical inspection, all devices were evaluated by leakage testing and impedance measurements of all electrodes. Per batch of 24, we subjected 3 devices to a thermal cycling stress test and confirmed leak tightness.

Gas-based leak testing^23^ was performed by sealing each device with an internal air pressure of 45 kPa, then measuring pressure decay over 30 s. For most devices, leakage was 0.006 kPa/s (**Figure 2f**). Rare leakages related to bubbles in the adhesive were detected by rapid pressure decay, occurring in <1% of devices.

Impedance measurements evaluated electrode functionality.^24^ Electrodes had a median impedance magnitude at 1 kHz of 47 kΩ (**Figure 2g**). Higher impedances of 100–250 kΩ may be related to contamination of the electrode surface; such electrodes are still functional but have higher noise.^24^ Less than 0.4% of electrodes with even higher impedances (often in the MΩ range) were assumed to be disconnected due to defects in the thin film traces or rivet bonding.

### Spheroid culture on mesh MEAs

We produced spheroids in 4 independent differentiations starting from neural progenitor cells,^25^ and placed spheroids on mesh MEAs after 3 or 5 weeks (**Figure 3**). In total, 239 spheroids were placed on individual mesh MEAs, of which we recorded from 157 (**Figure S2**). In this work, we used 89 mesh MEAs, with reuse up to 8 times (**Figure S3**).

**Figure 3.**
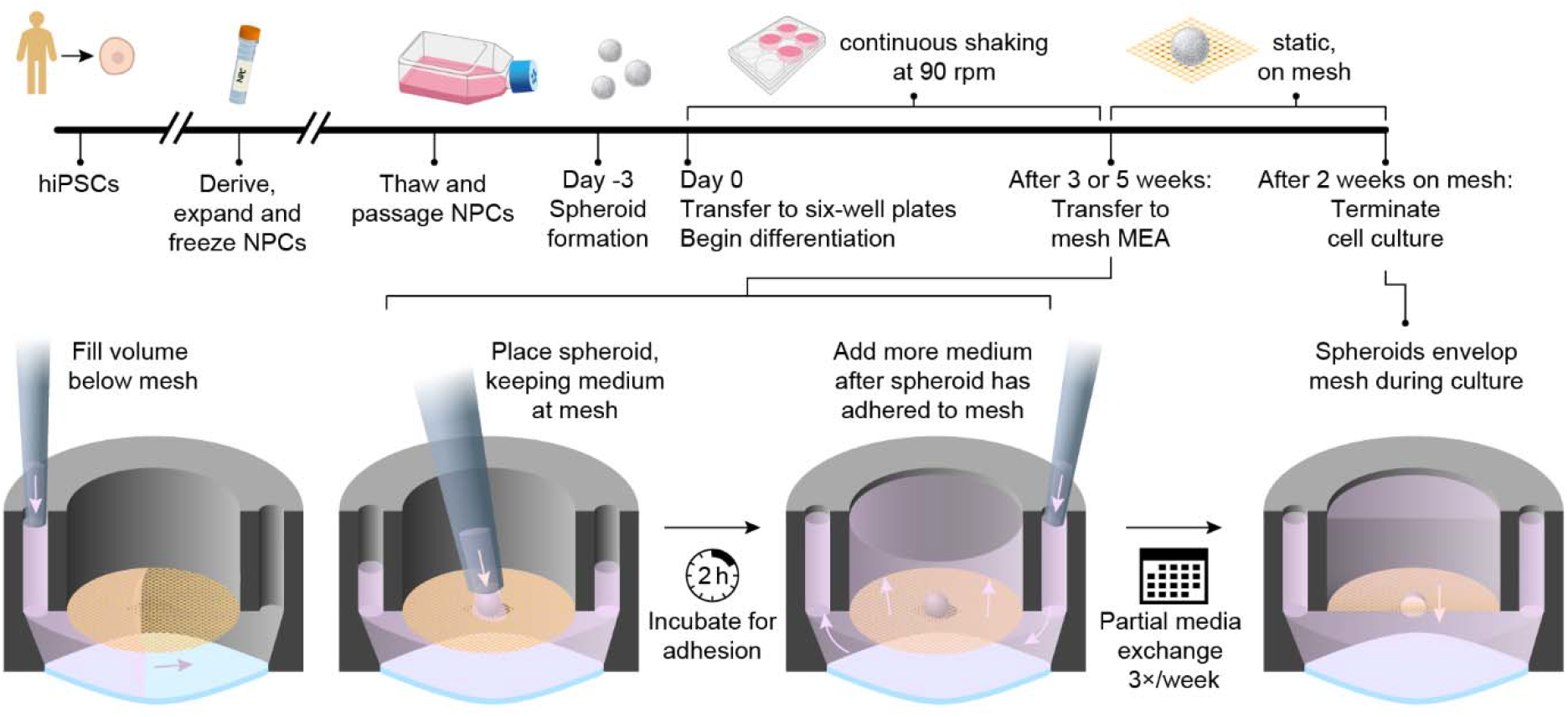
Spheroid generation, loading and culture on mesh MEAs. Above: timeline of spheroid generation and culture, before and on mesh MEAs. Below: Illustration of filling the device with medium, placing a spheroid on the mesh at the air– liquid interface for pinning until adhesion is established, then filling medium to a desired level.

Manual loading resulted in >95% of spheroids correctly placed on the central 1.4 mm square electrode field. Initially, maintaining medium level at the mesh immobilized spheroids. We used this air–liquid interface immobilization for 2 h, during which the spheroid adhered to the mesh. After achieving adhesion, adding medium did not displace the spheroids.

On the mesh, both imaging and electrophysiology track fine details of the spheroid structure. Over 2 weeks in culture on the mesh, spheroids maintained their round shape. Bright-field imaging revealed an increase in apparent size, as expected for these spheroids.^25^ Live/dead staining and confocal fluorescence imaging (**Figure 4**) further revealed that spheroids enveloped the mesh. The uptake of calcein AM by the first layers of cells revealed the morphology of the spheroids. Over 2 weeks of culture, spheroids sunk by up to 100 µm. This resulted in a slow insertion of electrodes into spheroids. We observed little interaction between cells and plasma-treated meshes, maintaining spheroid shape rather than dimpling or extending along mesh filaments. Importantly, we observed no overabundance of dead cells near the mesh. Another example is shown in **Figure S4** and **Video 1**.

**Figure 4.**
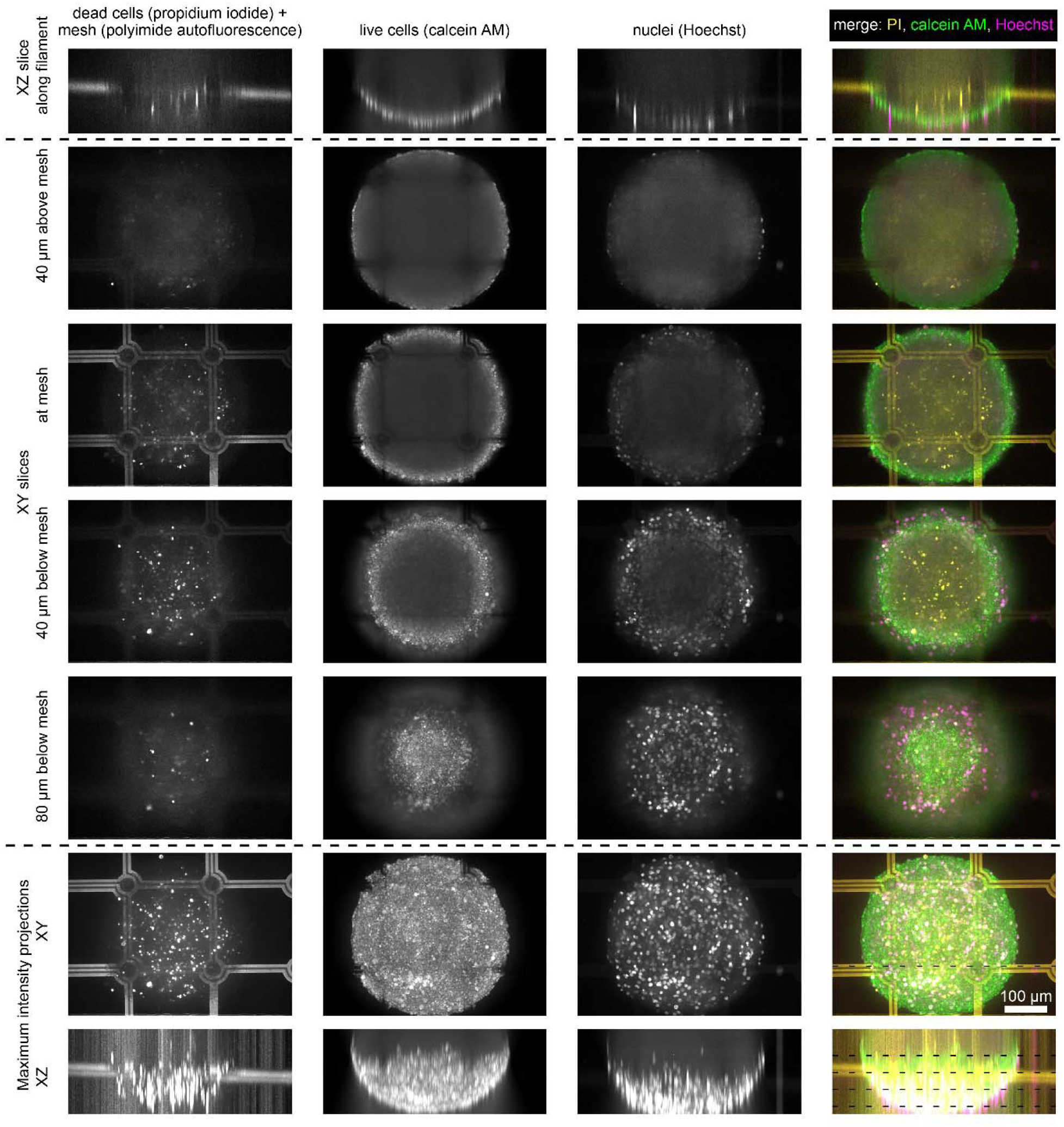
Neural spheroids adhere to and envelop mesh electrodes. This representative spheroid with a diameter of 300 µm sunk about 100 µm into the mesh over 2 weeks and then was stained for live and dead cells (calcein AM and propidium iodide, respectively) and nuclei (Hoechst). The spheroid maintained its round shape with minimal deformation at the mesh filaments, as shown by calcein fluorescence in the outer cells. Autofluorescence of the polyimide mesh appeared in the propidium iodide channel. We observed no overabundance of dead cells near mesh filaments. The location of orthogonal slices (above) is shown by dotted lines in the merged maximum intensity projection images (bottom right). The scale bar applies to all images.

### Electrophysiology

We recorded activity from spheroids from their second day on the mesh for up to two weeks. For each recording, mesh MEAs with spheroids were transferred from the incubator to the headstage, where temperature and gas environment were controlled. Spontaneous activity was recorded for up to 35 min to observe any instability due to transfer from the incubator to the headstage. In the first few minutes, we observed slow potential drift, however spiking activity stabilized quickly. For our analysis of each recording, we focused on a 5 min period at least 15 min after placing the MEA in the headstage.

Recordings of extracellular potentials revealed spiking activity with high fidelity (**Figure 5**). We identified spikes using threshold detection. Spontaneous activity was consistently observed over 2 weeks from electrodes in contact with spheroids, with spike rates in the range of 2–10 Hz. Synchronized bursts were observed across electrodes.

**Figure 5.**
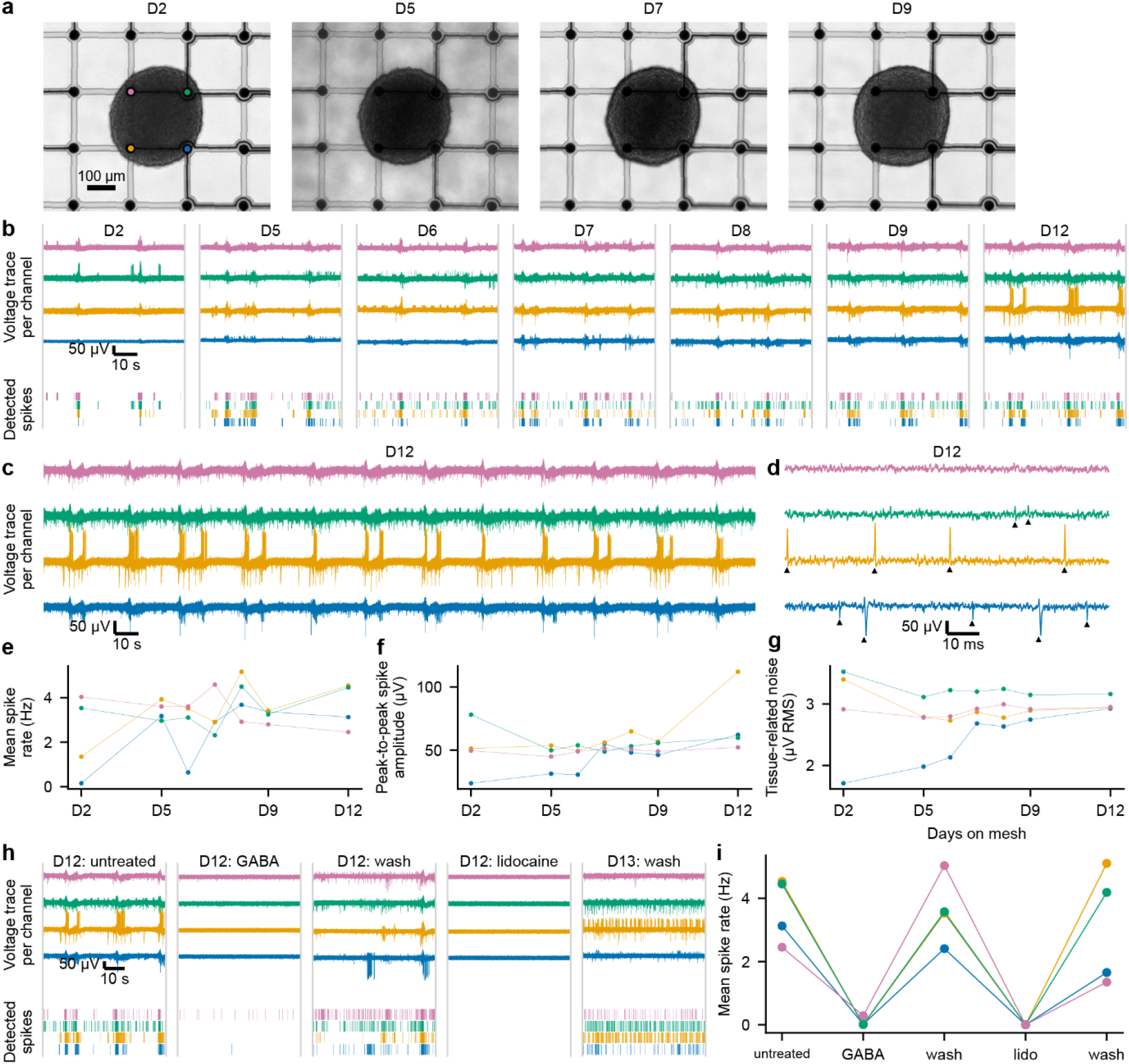
Mesh MEAs enable recordings from neural spheroids over many days, shown here for a representative spheroid. **a:** The spheroid remained round and no influence of the mesh (dimpling or bulging) was observed. Over 2 weeks, the apparent size of the spheroid increased. Colored electrodes in the first image indicate the electrode identity in the rest of the figure. **b:** Voltage traces and corresponding raster plots (below) from four electrodes, showing spontaneous activity including synchronized bursting. Recordings from a single spheroid, showing snapshots (1 min) from day 2 to day 9 on the mesh. **c:** Spontaneous activity over 5 min recorded on day 12 on the mesh, showing synchronized bursting, multiunit activity, and amplitudes >150 µV. **d:** A magnification from **c. e:** Mean spike rates over multiple days measured from each electrode were up to 5 Hz. The initial low rate on the blue electrode correlates with limited contact (see **g**). **f:** Amplitude of detected spikes over multiple days. **g:** Cell adhesion noise, calculated after high-pass filtering (3 kHz). Cells enveloped the blue electrode around day 7, although its proximity allowed it to detect signals before that (**e, f**). **h:** Pharmacological modulation produced expected effects, with inhibition by GABA or lidocaine. Each compound could be washed out, although residual effects of lidocaine reduced bursting activity even on the following day. **i:** Mean spike rates before, during and after pharmacological modulation (from **h**).

The example in **Figure 5** shows an interesting progression over 2 weeks. Synchronized bursting was observed in all recordings from 4 electrodes. Both the spike rate and amplitude varied as could be expected while the spheroid settles into the mesh. Although the spheroid appeared to cover 4 electrodes, closer examination of one electrode (blue in **Figure 5a**) suggests that it initially had no direct contact to the spheroid. On day 2 on the mesh, other electrodes appeared to have good contact and signals had higher amplitudes. On the blue electrode, signals had low amplitudes, mostly below the detection threshold (**Figure 5b**). Over the next 10 days, amplitudes at the blue electrode increased (**Figure 5e,f**). Notably, improved contact correlated with increasing apparent noise in the voltage traces. Thermal noise is related to cell adhesion,^26^ and high-pass filtering (3 kHz) to remove spiking activity confirmed that thermal noise increased as the blue electrode was enveloped (**Figure 5g**). The three other electrodes had good contact from day 2 and this was reflected in the cell adhesion noise. More distant electrodes (>100 µm from the spheroid) measured no activity and had low thermal noise.

After 12 days on the mesh, we added neurotransmitters or blockers to confirm the neuronal origin of the signals (**Figure 5h**). GABA strongly reduced spiking activity and could be readily washed out to restore activity. Lidocaine blocked all activity. Wash-out of lidocaine was slower, with activity measured the following day showing reduced bursting.

### Neural outgrowth via biofunctionalization

Laminin coating of the mesh promoted migration and outgrowth of cells from the spheroid (**Figure 6**). This behavior contrasted starkly with the plasma-treated, uncoated meshes (**Figure 6e**). On uncoated meshes, recordings revealed activity typically limited to four electrodes. On laminin-coated meshes, activity was reliably measured as cells spread to most electrodes over two weeks. Outgrowth along mesh filaments was observed by bright-field imaging. Fluorescent staining revealed both neurite outgrowth and soma migration. Impedance spectroscopy correlated with outgrowth (**Figure S5**), supporting its use for label-free monitoring in 3D cellular assays.

**Figure 6.**
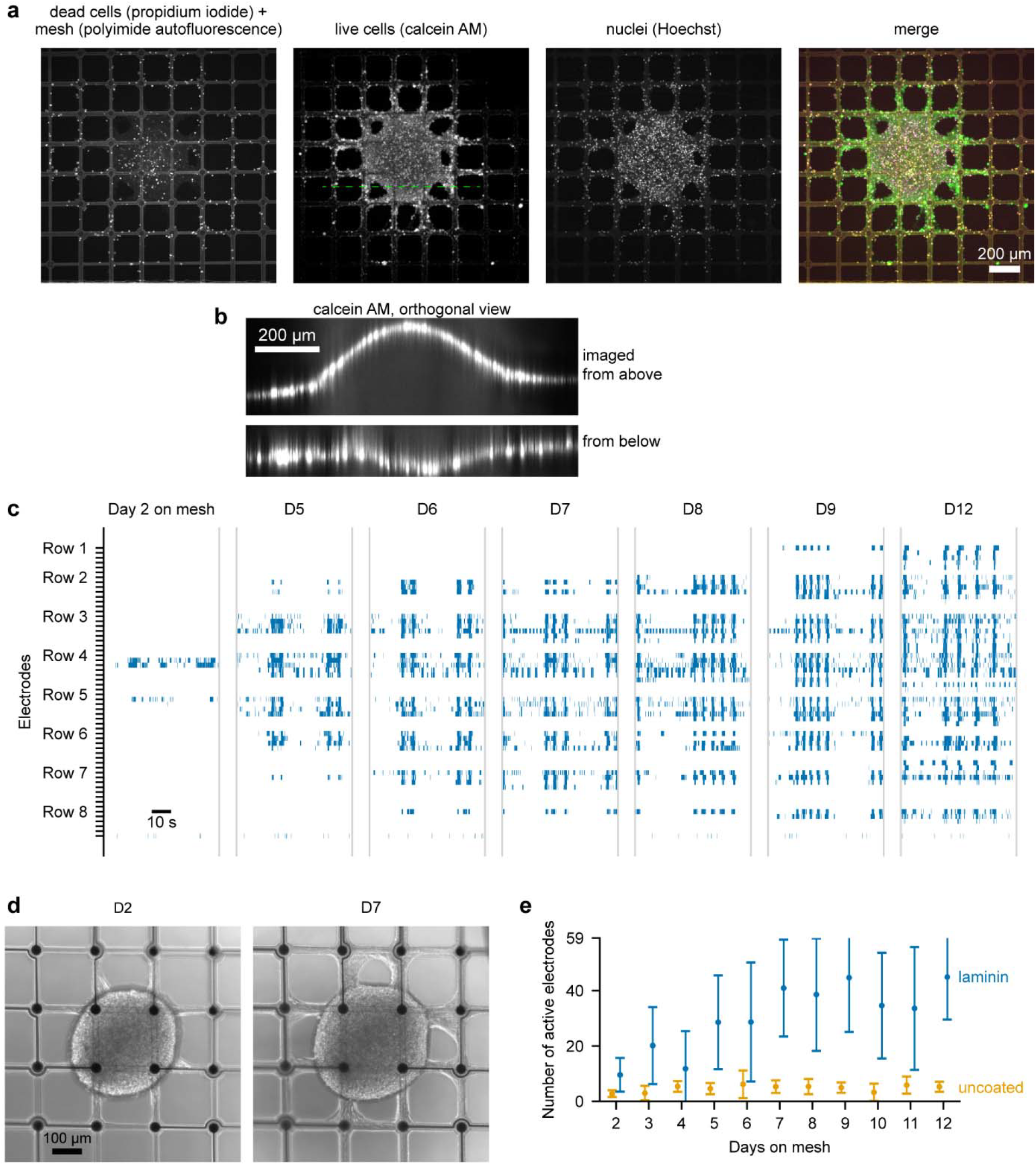
On laminin-coated meshes, cells migrate and extend neurites along the filaments. Electrical activity matches the observed spreading. **a:** A representative spheroid after 2 weeks on the mesh, stained for live and dead cells (calcein AM and propidium iodide, respectively) and nuclei (Hoechst). **b:** Orthoganol projections revealed that the spheroid deformed as it spread across the mesh, as shown by calcein fluorescence in its outer cells. The green dashed line in **a** indicates the location of orthogonal slices. **c:** A representative raster plot shows how spontaneous activity spread to the majority of the electrodes after a few days on the mesh. **d:** Bright field images of the spheroid from **c. e:** Spreading of cells on laminin-coated meshes rapidly increased the number of active electrodes. In comparison, spheroids on uncoated meshes maintained a low number of active electrodes over time.

## Discussion

This study demonstrated that a suspended, topologically 2D mesh MEA can provide longitudinal extracellular recordings from within neural spheroids while preserving their overall morphology. The method combines gradual tissue integration, pharmacologically responsive electrophysiological readouts, and practical handling within a manufacturable device format. Our design deliberately prioritized experimental robustness, manufacturability, and compatibility with established MEA workflows over maximal mechanical conformability or electrode density. In vitro models, beyond recreating physiological features, must consider challenges of experimental variability, numbers of samples, experimental complexity, and cost.^19^ Success will not be the most advanced technology, but rather better understanding of human physiology and the development of therapeutics.

Mesh MEAs diverge from conventional MEAs based on solid planar substrates to better fit to 3D in vitro models. Although microfabrication begins on rigid substrates, meshes are then detached and assembled into final devices. Devices range from our 2D mesh format, to various flexible meshes for organoids^14,16^ or animals,^27^ or even 3D MEAs using shanks^28,29^ or stacked meshes^30,31^. These approaches occupy different design spaces rather than a single hierarchy of performance.^10^ More conformable or fully three-dimensional devices can provide different forms of tissue coverage or spatial sampling, but often require more complex fabrication, handling, implantation, or model-specific adaptation. Our approach is complementary: it maintains a fixed electrode geometry and standard MEA-like handling while still allowing cells to envelop the mesh and bring electrodes into the spheroid.

Reducing production effort required microfabrication of mesh chips on batches of carrier wafers, followed by semiautomatic wire bonding to EIBs.^32^ A compact well design enabled simple assembly and ease of use.^33^ This philosophy has guided development over multiple device generations focused on reliability, usability, and manufacturability (**Figure S6**). Generation 3 has proven to be reliable (up to 99.6% functional electrodes, **Figure 2g**) and even reusable (**Figure S3**). A fourth generation is in development, following design-for-manufacturing principles for simpler assembly and automated adhesive bonding to reduce production costs.

We were conservative in choosing materials. The mesh is made of polyimide, which has been demonstrated to be biocompatible and stable in implants,^34^ and together with TiN electrodes and titanium/gold traces is well established in MEA technology.^11,15^ The well is made of polycarbonate, and we avoided the risk of cytotoxicity from 3D printed polymers.^35^ We used an epoxy adhesive which has proven to be stable and biocompatible in previous work.^7,15,36^ Copper in the EIB could corrode if in contact with salt water, however it is protected with a gold surface. The gold bonds between the mesh MEA and EIB were chosen to avoid risks of corrosion.

Beyond biocompatibility, our conservative material choices also enabled reusability, which is an important feature for MEAs. We have not studied reusability systematically, but rather reused mesh MEAs as needed, up to 8 times (**Figure S3**). Each use involved sterilization in ethanol, 2 weeks in culture, and cleaning. Signal quality remained high, and no adhesive failure or leakage was observed during cell culture. Further testing is required to understand the real limitations of mesh MEAs.

Optimized manual liquid handling achieved a >95% success rate in placing spheroids on the electrode field (**Figure S7**). In contrast, preliminary results had a 25% success rate: of 24 spheroids placed on the electrodes, 18 were displaced during transport from the sterile bench to the incubator. The key improvement was to pin the spheroid in place using the air–liquid interface at the mesh for 2 h after placement. After 2 h, medium was added and we observed no movement of spheroids. Longer air–liquid interface culture is possible but must consider the low 100 µL volume below the mesh.

The excellent adhesion of spheroids to the mesh is an important feature. Monolayer cultures typically require coatings on MEAs to ensure adhesion of cells. Spheroids are more difficult, easily rolling away and preventing stable recordings. We observed good adhesion on mesh MEAs, even without any coating.

Spheroids, once adhered, showed no movement over 2 weeks, including regular medium exchange and recordings.

The electrophysiological recordings indicate that gradual mesh envelopment translated into functional electrode–tissue coupling. Increasing spike amplitudes and adhesion-related noise were consistent with improved cellular contact, while pharmacological suppression by GABA and lidocaine supported the neuronal origin of the recorded signals. The recordings further captured synchronized bursting across electrodes, demonstrating that the device reports not only local spiking activity but also network-level dynamics within the spheroid. At the same time, the present analysis remains based on threshold-detected extracellular events and does not yet assign individual waveforms to defined neuronal subtypes or putative single cells. Future work should combine spike sorting, immunocytochemistry, and optical correlation to relate recorded waveforms to local cytoarchitecture and specific neuronal populations.

Interactions of cells with the mesh, including adhesion, motility, and envelopment, require further investigation. Our results suggest that mesh surface chemistry can define two distinct modes of tissue interaction. On native polyimide meshes, after plasma treatment but without protein coating, spheroids maintained their compact morphology while gradually enveloping the mesh. This represents a spheroid-preserving mode, in which electrodes become embedded without overt disruption of the overall tissue shape. We suspect that dynamic intercellular adhesion may cause spheroids to envelop the mesh, perhaps supported by gravity due to the higher density of spheroids in comparison to culture medium. A laminin coating changed the mode of tissue–mesh interaction, causing cell spreading, neurite outgrowth, and soma migration, which reduced spheroid settling while increasing electrode coverage (**Figure 6**). This represents a bioengineering mode, in which surface functionalization can be used to guide cell migration and network extension along the mesh. Different functionalization strategies may therefore pursue either a low-interaction, spheroid-preserving interface or an instructive interface for engineered models, for example by using patterned, cell-type-specific adhesive molecules to guide connections between assembloids.

The cytoarchitecture near the mesh will also be interesting to study. As spheroids envelop the mesh, local neurite geometry may be altered around mesh filaments, and cells may need to remodel their processes at the tissue–device interface. Of course, neurites can be dynamic, even growing by hundreds of micrometers per day.^37^ Such effects will depend on the maturity and biomechanical properties of a spheroid. Organoids with well-defined layering may benefit from mesh layouts with fewer filaments but more closely spaced electrodes. More challenges may be faced by spheroids with stronger intercellular connections, such as heart or muscle organoids.

We did observe technical challenges when using these mesh MEAs. Unintentional contact of a pipette tip to the mesh easily tore it, creating a hole and breaking electrode traces. In one case, a mesh MEA device broke during removal of the membrane chamber. We attribute this to friction against the vertical outer wall of the well; the outer wall is tapered in our generation 4 design. During recordings, we observed transient session-specific noise on some recording channels, while the same channels recorded activity with low noise on other days. In contrast, a small number of defective electrodes always recorded high noise. We attribute transient problems to poor electrical contact between the headstage’s gold pins and the EIB pads, despite cleaning with isopropanol before each recording. Interconnection reliability will be important for mesh MEAs with higher numbers of channels or in multiwell formats.

## Methods

### Device design

We designed mesh chips in CleWin 5 (WieWeb, the Netherlands). The mesh was defined as a square grid with 200 µm pitch over a circular region with a diameter of 7 mm. Electrodes (diameter 30 µm) were positioned at the central 8×8 nodes and traces were routed along filaments to the mesh circumference.

Routing used a maximum of 2 traces along any filament. Filaments were 21 µm wide.

We designed EIBs as printed circuit boards (PCBs) using KiCaD. EIBs had two layers with 90 µm lines and spaces, 35 µm copper, bonding gold (ENEPIG) finish, and a high temperature resin (FR4, HTg 180) with 1 mm thickness.

Well parts were designed in SolidWorks and were CNC milled from polycarbonate. Bottom windows were laser-cut from polycarbonate foil.

### Microfabrication of mesh chips

We produced mesh chips on 100 mm silicon carrier wafers.^32,33^ The layer stack was polyimide 1 (6 µm), titanium/gold/titanium (TiAuTi, 0.5 µm), polyimide 2 (6 µm) and titanium nitride (TiN, 0.5 µm). Three photolithography processes were used to pattern the traces, to structure polyimide and open the electrodes, and to pattern the electrode material. The mesh chips were visually inspected under a microscope for defects before being removed from the wafer using tweezers. Detached mesh chips were stored between layers of cleanroom paper.

### Wire bonding

EIBs were produced by a commercial PCB supplier and first cleaned with isopropanol. Single mesh chips were placed onto the EIB and manually aligned to the bond pads. Alignment was facilitated by a drop of distilled water. Evaporation of water temporarily fixed the mesh chip to the EIB. Bond contacts used through-holes surrounded by a gold ring (**Figure 2e**), and each contact had two bond sites for redundancy. Rivet bonding^20^ connected the mesh contacts via their through-holes to the EIB. Rivet bonding used a semiautomatic F&K Delvotec 5610 wire bonder, a 38 µm gold wire and a UTS-51JM-AZ-1/16-16MM tip (SPT Roth Ltd). Parameters were: ball size: 5.0, flame off current: 40 mA and 10 kΩ, tail height: 40 µm, bond force: 40 cN, ultrasonic time 38 ms, ultrasonic power: 80 digit.

### Assembly and adhesive bonding

Well parts were cleaned in isopropanol with ultrasonication for 5 min before assembly. Prior to adhesive bonding, all parts were treated with air plasma (90 s, 150 W, 0.5 mbar, Pico from Diener electronic). EPO-TEK 301-2FL was prepared in a ratio of 100:35 according to manufacturer’s guidelines and mixed for min at 3000 rpm using a SpeedMixer (Hauschild Engineering). Possible crystals in part A were melted by heating in a water bath no longer than 2 weeks before use (and cooled to room temperature before use). Wells and meshes (rivet bonded to EIBs) were assembled manually. The well top and bottom sandwiched the mesh. Alignment was achieved by posts on the bottom part, which passed through holes in the mesh (**Figure 2c**) into holes in the top part. A window was placed below the well bottom. Before applying adhesive, all parts were clamped together using 3D printed jigs. Adhesive was applied manually using a Nordson dispenser (Ultimus I, tips with an inner/outer diameter of 0.1/0.24 mm, 100–150 kPa) at the mesh/polycarbonate interface and at the window. While clamped, the adhesive was cured at 80°C for 3 h. After cooling, jigs were removed.

A second gluing step added so-called well expanders, on top of the 9 mm-square well tops. Partially assembled mesh MEAs and the well expanders were plasma treated and the adhesive was prepared as above. Well expanders were positioned requiring no clamping, and adhesive was dispensed and again cured at 80°C for 3 h.

### Impedance measurements

Mesh MEAs were treated with air plasma then filled with phosphate-buffered saline (PBS). Impedance magnitude at 1 kHz was measured using a MEA-IT system (Multi Channel Systems MCS GmbH, Germany) and an external Ag/AgCl reference electrode. Electrodes having impedance greater than 500 kΩ at 1 kHz were classified as defective. Impedance spectra were measured using a Gamry 600+ potentiostat.

### Leakage testing, hygrothermal aging and thermal cycling

Leak tightness was tested by sealing each device with positive air pressure (40–50 kPa) and measuring pressure decay. Each mesh MEA was sealed against a rubber gasket. Pressure was applied, the inlet was closed, and decay was measured for 30 s. A discretionary threshold was used to differentiate sealed and leaky devices. The threshold was defined as 0.25% of the initial pressure per second (0.11 kPa/s at 45 kPa).

Per assembled batch of mesh MEA devices, representative devices were sterilized in ethanol for 30 min, filled with PBS and subjected to a 5-day thermal cycling process. Thermal cycling used hourly cycles between 10°C and 47°C at 98% relative humidity in a Vötsch (VCL 4010) climate chamber. Subsequent leakage testing was performed for batch release.

### Spheroid generation and culture

Each differentiation of neural spheroids (**Figure 3**) began with thawing of neural progenitor cells (NPCs). NPCs were derived from the KOLF2.1J iPSC line (JIPSC1000, The Jackson Laboratory, CT, USA) following Reinhardt et al., 2013^38^ before freezing. Culture plates were coated with 5 µg/mL laminin (A29248, Thermo Fisher) in DPBS +/+ overnight at 4°C. NPCs were plated at a concentration of 50,000 cells/cm^2^ and cultured in NPC culture medium (**Table 1**) at 37°C and 5% CO_2_. Medium was refreshed three times per week until 80% confluency was reached.

**Table 1.** NPC culture medium composition. In specified steps, the medium was used without growth factors BDNF, FGF and EGF.

| Item | Concentration | Product info |
| --- | --- | --- |
| Neurobasal Medium | 1:1 with DMEM/F12 medium | 21103049, Gibco, Thermo Fisher Scientific, MA, USA |
| DMEM/F-12, GlutaMAX supplement | 1:1 with Neurobasal medium | 31331028, Gibco, Thermo Fisher Scientific, MA, USA |
| B-27 Supplement (50X), minus vitamin A | 1X | 12587010, Gibco, Thermo Fisher Scientific, MA, USA |
| N-2 Supplement (100X) | 1X | 17502048, Gibco, Thermo Fisher Scientific, MA, USA |
| Penicillin-Streptomycin (10,000 U/mL) | 100 U/mL | 15140122, Gibco, Thermo Fisher Scientific, MA, USA |
| 2-mercaptoethanol (50 mM) | 50 µM | 31350010, Gibco, Thermo Fisher Scientific, MA, USA |
| Human/Mouse/Rat BDNF, Animal-Free Recombinant Protein, PeproTech® | 20 ng/mL | AF-450-02, Gibco, Thermo Fisher Scientific, MA, USA |
| Human FGF-basic (FGF-2/bFGF) (154 aa), Animal-Free Recombinant Protein, PeproTech® | 10 ng/mL | AF-100-18B, Gibco, Thermo Fisher Scientific, MA, USA |
| Recombinant Human EGF Protein, CF | 10 ng/mL | 236-EG, R&D Systems Inc., MN, USA |

After reaching 80% confluency, NPCs were detached by incubation with StemPro Accutase Cell Dissociation Reagent (A1110501, Gibco, Thermo Fisher) for 5 min at 37°C with 5% CO_2_. Once the cells were detached, Accutase was neutralized with the NPC culture medium without growth factors, and the cells were centrifuged at 300×g for 5 min.

In preparation for aggregation of NPCs into spheroids, 24-well AggreWell 800 culture plates (34815, STEMCELL Technologies) were treated with anti-adherence rinsing solution (07010, STEMCELL Technologies). Each well was filled with 500 µL of solution, and plates were centrifuged at 1300×g for 5 min. The wells were then rinsed twice with NPC culture medium without growth factors (**Table 1**).

Detached NPCs were resuspended in NPC culture medium and plated in anti-adherent AggreWell 800 culture plates at a concentration of 1.5 x 10^6^ cells per well, resulting in 5000 cells per spheroid. Plates were centrifuged at 100×g for 3 min. After confirming uniform cell distribution, plates were incubated at 37°C with 5% CO_2_. After 3 days, medium in the AggreWell 800 was replaced with spheroid differentiation medium (**Table 2**). Neural spheroids from each well were then transferred to individual wells of a 6-well plate. Each well was supplied with 2 mL of spheroid differentiation medium, and 75% of medium was exchanged three times per week. Plates were incubated at 37°C with 5% CO_2_ on an orbital shaker (90 rpm, 19 mm orbit diameter, Thermo Scientific CO_2_ Resistant Shaker). After 3 weeks in spheroid differentiation medium, cultures were switched to spheroid maturation medium (**Table 3**). Per differentiation, spheroids were transferred to mesh MEAs at 3 weeks or 5 weeks (**Figure S2**).

**Table 2.**
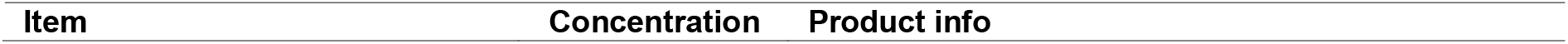
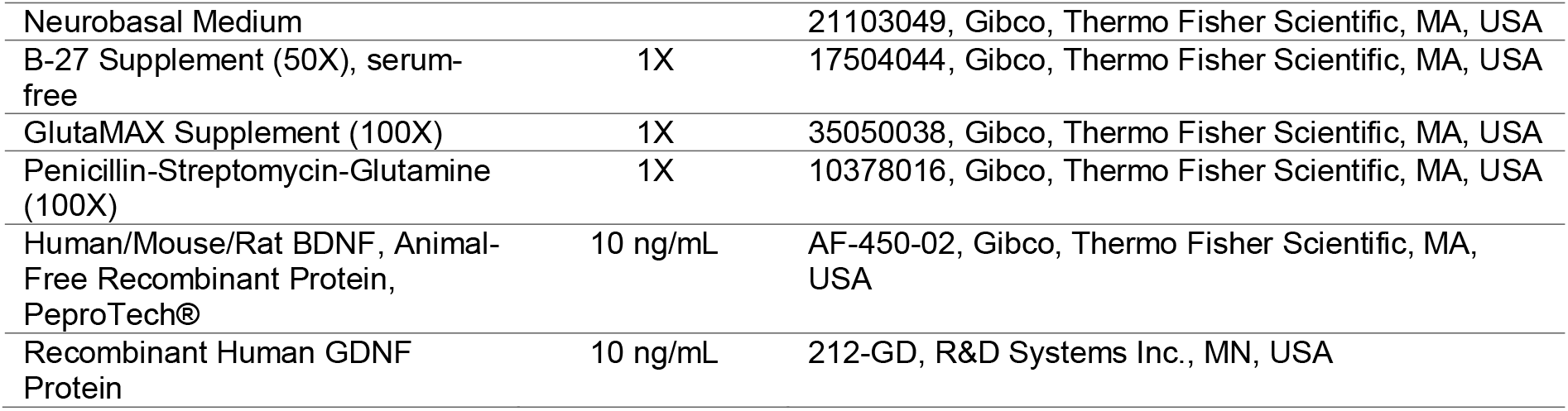
Spheroid differentiation medium composition.

| Item | Concentration | Product info |
| --- | --- | --- |
| Neurobasal Medium |  | 21103049, Gibco, Thermo Fisher Scientific, MA, USA |
| B-27 Supplement (50X), serum-free | 1X | 17504044, Gibco, Thermo Fisher Scientific, MA, USA |
| GlutaMAX Supplement (100X) | 1X | 35050038, Gibco, Thermo Fisher Scientific, MA, USA |
| Penicillin-Streptomycin-Glutamine (100X) | 1X | 10378016, Gibco, Thermo Fisher Scientific, MA, USA |
| Human/Mouse/Rat BDNF, Animal-Free Recombinant Protein, PeproTech® | 10 ng/mL | AF-450-02, Gibco, Thermo Fisher Scientific, MA, USA |
| Recombinant Human GDNF Protein | 10 ng/mL | 212-GD, R&D Systems Inc., MN, USA |

**Table 3.** Spheroid maturation medium composition.

| Item | Concentration | Product info |
| --- | --- | --- |
| BrainPhys Medium |  | 5790, STEMCELL Technologies Inc., Germany |
| B-27 Supplement (50X), serum-free | 1X | 17504044, Gibco, Thermo Fisher Scientific, MA, USA |
| GlutaMAX Supplement (100X) | 1X | 35050038, Gibco, Thermo Fisher Scientific, MA, USA |
| Penicillin-Streptomycin-Glutamine (100X) | 1X | 10378016, Gibco, Thermo Fisher Scientific, MA, USA |
| Human/Mouse/Rat BDNF, Animal-Free Recombinant Protein, PeproTech® | 10 ng/mL | AF-450-02, Gibco, Thermo Fisher Scientific, MA, USA |
| Recombinant Human GDNF Protein | 10 ng/mL | 212-GD, R&D Systems Inc., MN, USA |

### Loading spheroids on mesh MEAs

Before loading spheroids, mesh MEAs were treated with low pressure air plasma for 90 s at 150 W. They were then moved to a sterile bench and disinfected by immersion in 80% ethanol for 30 min. Ethanol was removed and devices were allowed to dry.

For laminin coating, the mesh MEAs were filled with 50 µg/ml poly-L-ornithine (PLO, Advanced Biomatrix) and put in the incubator at 37°C and 5% CO_2_ overnight. The next day PLO was removed, and each mesh MEA was rinsed three times with PBS before 1 µg/ml laminin solution (STEMCELL Technologies) was added. The mesh MEAs were put back in the incubator for at least 1 h. Laminin solution was removed and the mesh MEAs were rinsed three times with PBS, then were used directly or refrigerated filled with PBS to use the next day.

For spheroid loading (**Figure 3**), the volume below the mesh was filled with 100 µL of medium. For each mesh MEA, a petri dish was prepared with a filter paper with a central viewing hole, wetted with 1 mL deionized water for humidity control. Using a 200 µL pipette tip set to 6 µL and working under a microscope, individual spheroids were picked from a 6-well plate and placed onto the electrode field. Any medium above the mesh was removed, pinning the spheroid in place by surface tension. After loading, MEAs were placed in an incubator for 2 h to allow pinned spheroids to adhere to the mesh. Then, 150 µL medium was added to the inner well.

Spheroids from 4 differentiations were loaded on mesh MEAs. Per differentiation, at least 24 spheroids were loaded on individual mesh MEAs, and a total of 157 spheroids were recorded (**Figure S2**).

### Spheroid culture on mesh MEAs

Medium was partially exchanged (60%/150 µL) three times a week on Monday, Wednesday and Friday. Spheroids were imaged on the day of medium exchange, inside the petri dish using an inverted bright field microscope (**Figure 5a**).

Spontaneous activity was recorded at least 3 times per week for 2 weeks. Pharmacological experiments and live/dead staining were performed after 2 weeks on the mesh.

### Electrophysiology

Spontaneous electrical activity was recorded at least 3 h after medium exchange. Prior to recording, mesh MEAs were removed from their petri dishes and covered with membrane chambers (MEA-MEM, MCS).

Gold contact pads and gold pins of the amplifier were cleaned using isopropanol and a cotton swab before connecting. The heating stage was set to 37°C, and the MEA was supplied with 95% O_2_, 5% CO_2_ gas flow, humidified by flowing through a bubbler. Electrical activity was recorded using a MEA2100 amplifier (Multi Channel Systems MCS GmbH), with sampling at 25 kHz after filtering (10 Hz high-pass, 7 kHz low-pass, second-order Butterworth filter).

### Pharmacology

Stock solutions were prepared and stored at 4°C. Solutions in medium were prepared on the day of the experiment, and diluted to a final concentration in the device. Concentrations (stock, final) were 10 mM, 100 µM (GABA); 100 mM, 1 mM (lidocaine); and 100 mM, 1 mM (CdCl_2_).

Baseline recordings were first acquired. For compound application, the membrane chamber was removed, and 20 µL of medium was replaced and mixed in the device, and the chamber was replaced. After 5– 10 min, activity was recorded for 5 min. Afterwards, medium was fully exchanged (200 µL) and MEAs were returned to the incubator. Recordings after washing were repeated on the same day or the following day for lidocaine.

### Imaging

Staining solution was prepared by mixing 4 µL calcein AM, 4 µL Hoechst 33342 and 50 µL propidium iodide. Spheroids were incubated for 30 min with 4.8 µL staining solution diluted in 200 µL medium in the mesh MEA. Confocal imaging used a Cell Observer (Zeiss). Excitation and detection filters were 509 nm and 488 nm (calcein AM), 455 nm and 352 nm (Hoechst), and 561 nm and 548 nm (propidium iodide).

### Device cleaning

After use, 2 mL of Tergazyme solution (Alconox, Inc.) was added to the mesh MEA and left overnight. Then all liquid was removed, mesh MEAs were washed with distilled water and left to dry. MEAs were inspected for biological residue and physical damage before reuse.

### Data analysis

Electrical recordings were analyzed using custom python scripts. For spike detection, raw signals were filtered (0.3–3 kHz second-order Butterworth band-pass) and events were identified by threshold detection (mean ± 6 × standard deviation). Any data points within 3 ms were reduced to a single event.

## Supporting information

Confocal scan through a spheroid on a mesh MEA

## Acknowledgements

We thank our collaborators for their valuable feedback on early prototypes. We thank Thoralf Herrmann for EIB design, Stefan Klaus for cleanroom support, and Elvina Houas and Roswitha Fischer for assembly and inspection support. We thank Sven Schönecker, Jannis Meents and Moritz Gerlach from MCS for discussions about quality control. We thank Lisa Brauns from NMI TT GmbH for discussions about manufacturability and quality control. T.S. and P.D.J acknowledge funding by the German Research Foundation (DFG) in project MEMMEA (441918103) as part of the DFG priority program SPP 2262 MemrisTec (422738993). P.L. and L.E. were supported by the Innovative Medicine Initiative 2 Joint Undertaking (JU) under grant agreement no. 853988. The JU receives support from the European Union’s Horizon 2020 research and innovation program and EFPIA and JDRF International. This work received financial support from the State Ministry of Baden-Wuerttemberg for Economic Affairs, Labour and Tourism. T.S., M.M. and P.D.J. are inventors on a patent application related to mesh MEAs.

