## Supplementary material for "A scalable mesh microelectrode array platform for longitudinal electrophysiology in neural spheroids"

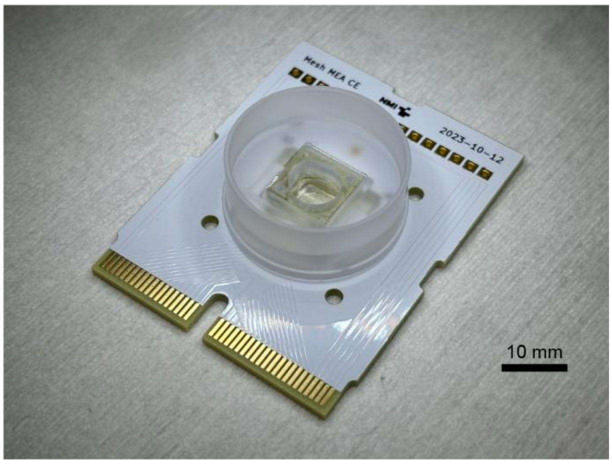


**Figure S1:** Mesh MEA using a card-edge EIB compatible with 80-channel Samtec HSEC8 connectors. Keeping all contacts along one edge simplifies fluidic interfacing. For example, the EIB can be connected without disconnecting any tubing connections.

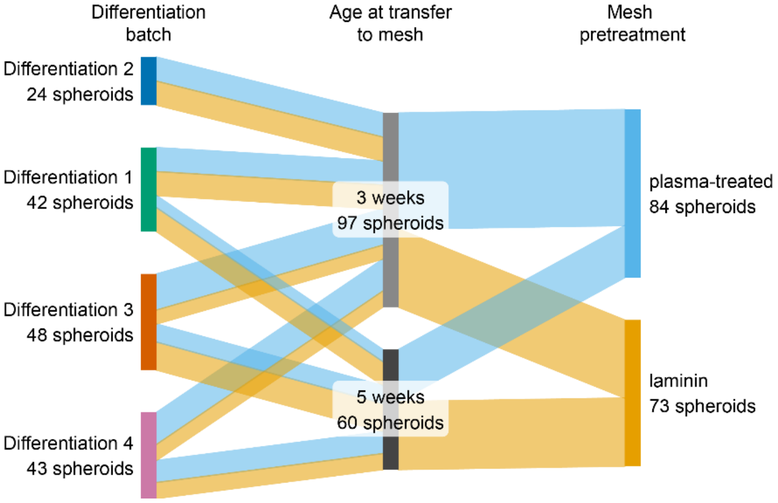


**Figure S2:** Sankey diagram of 157 spheroids from 4 differentiations transferred to and recorded using mesh MEAs. Spheroids from differentiation 2 intended for transfer to mesh MEAs at week 5 were not used due to a logistical error. In each differentiation, reserve spheroids were transferred to mesh MEAs but not recorded; these are excluded here.

*

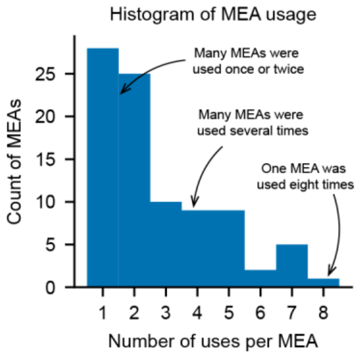


**Figure S3:** Mesh MEAs can be reused after cleaning. Most mesh MEAs in this work were used once or twice, while several were used 6–8 times.

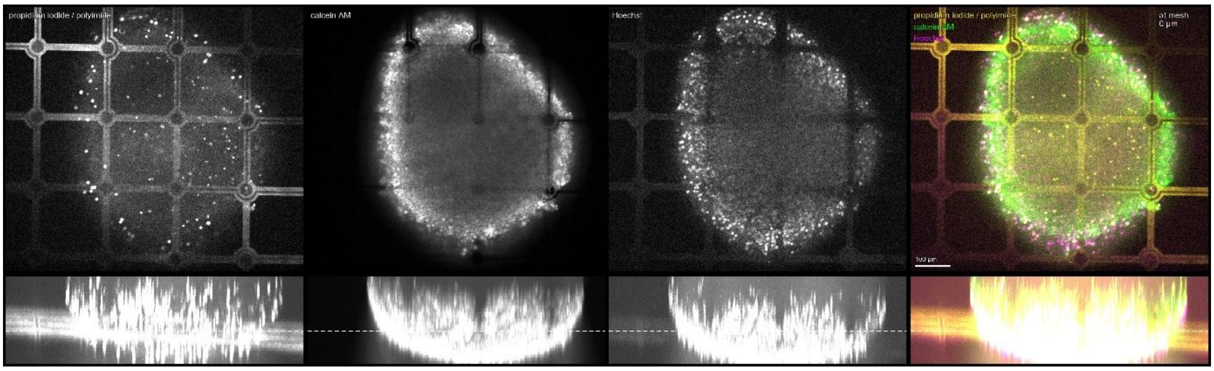


**Figure S4:** Snapshot of video scanning vertically through a confocal stack. Electrodes were 100 µm into the spheroid. The vertical position of the xy slice is indicated by a dashed white line on the orthogonal maximum intensity projection. For details on staining, refer to **Figure 4**.

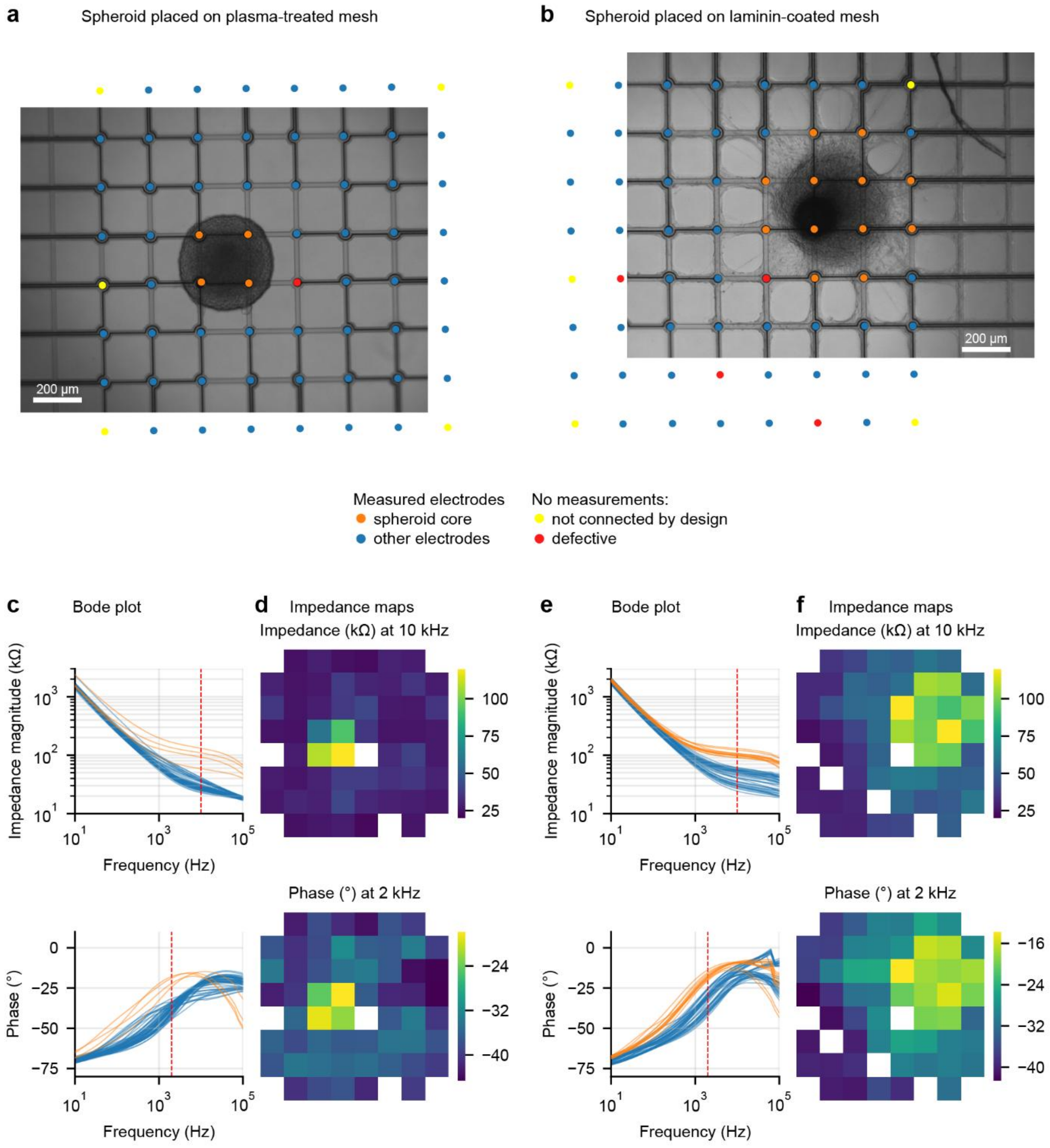


**Figure S5:** Impedance spectroscopy reveals presence of tissue for a round spheroid on a plasma-treated mesh (**a**) and a spread-out spheroid on a laminin-coated mesh (**b**), both after 2 weeks of culture on the mesh. Impedance spectra (**c**, **e**) show a clear difference between electrodes in the spheroid core (orange) vs. other electrodes (blue). Mapping of impedance or phase at single frequencies (**d**, **f**) matches the observed presence of cells.

### Four generations of scalable mesh MEAs

The device generation (1, 2, 3, 4) specifies the well construct and its assembly process, but not the EIB. For example, generation 3 devices were produced using the 49×49 mm² 59-channel EIB for MCS amplifiers but also a 64-channel card-edge EIB (**Figure S1**). In all cases, we've used the same mesh design with a 200 µm filament pitch and 8×8 microelectrodes. However, the generation is agnostic to the mesh design, as long as it fits in the 7 mm inner well.

In the first generation, some devices were stable during culture for several weeks, while others developed leaks. The design of the well was sensitive to the manual assembly process: too much adhesive risked blocking inlets or contaminating electrodes, while too little resulted in poor stability.

In generation 2, we improved adhesive bonding by including capillary structures near the bonding interface. With the parts clamped together, the capillary structures defined the adhesive bondline thickness and provided a reservoir to distribute adhesive while avoiding overflow. The first generation had a glass window at the bottom of the well. Despite functionalizing glass using an aminosilane adhesion promoter, thermal stress prevented a reliable glass–polymer interface. Replacing the glass window with polycarbonate resolved this issue, and neither noticeably increased autofluorescence nor degraded optical clarity. A further process improvement was to heat the epoxy resin to eliminate crystals, which can prevent complete mixing and curing (EPO-TEK Tech Tip 7).

In generation 3, we increased the well size, adding the larger well with compatibility for membrane caps. The well size allows larger medium volume, although we used low medium volumes within the inner well. The membrane caps were important for reducing risks of contamination and evaporation during recording in headstages outside of the incubator. This design was used for all validation in this paper.

The fourth generation maintains the materials of generation 3. Changes to user-facing features include adding a slope to the walls of the outer ring and decreasing the height of the inner 7 mm-diameter well above the mesh. The electrical connections between the mesh and the EIB, as well as the components to form the well were redesigned to improve manufacturability, so that critical assembly steps may be automated.

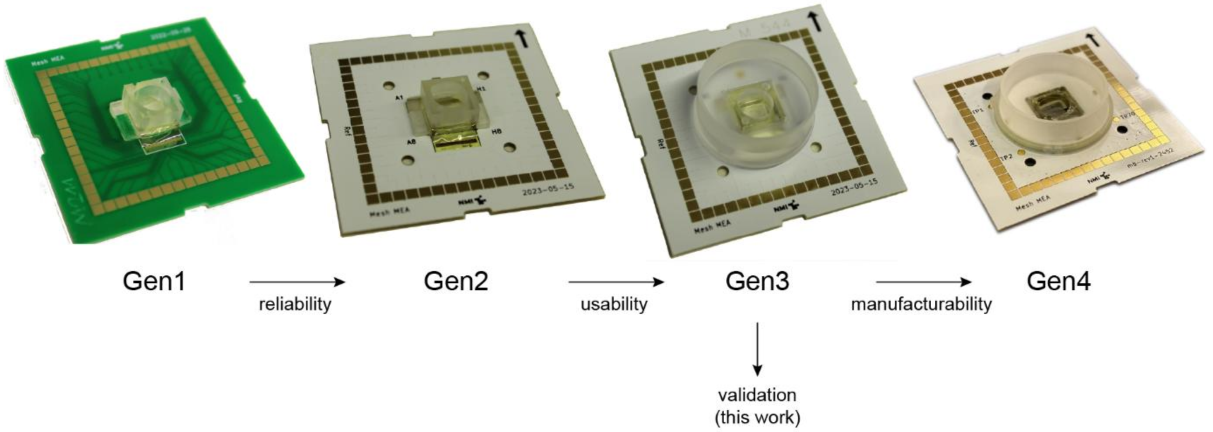


**Figure S6:** Overview of mesh MEA device generations.

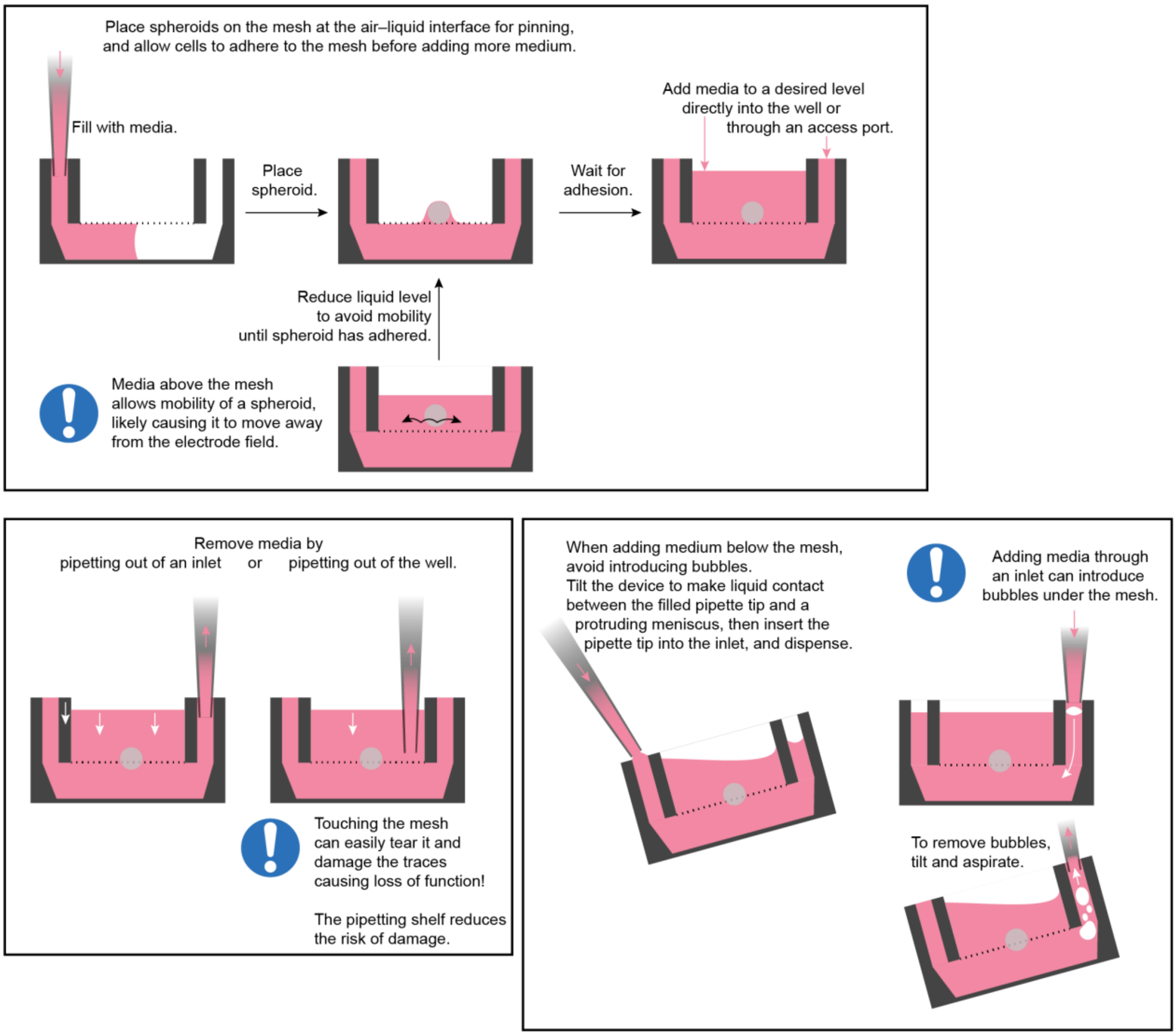


**Figure S7:** Schematics for liquid handling in mesh MEAs. **a:** After placing a spheroid, pinning for 2 h using the air–liquid interface resulted in good adhesion of the spheroid on the mesh. **b:** Removing medium required care to avoid damage to the mesh. Medium should be preferably removed from an access port. If medium must be removed from the inner well directly, the pipette tip can be placed on the pipetting shelf (**Figure 1c**). **c:** Care is required to avoid injecting bubbles below the mesh. Bubbles can be removed by tilting towards an inlet and aspirating the bubbles.
